# Constitutive sporulation in wild fission yeast enhances insect-mediated survival

**DOI:** 10.64898/2026.01.14.699589

**Authors:** Hironori Niki, Shunsuke Noda, Keigo Fujiwara, Takuya Miyake, Koichiro Akiyama, Taisuke Seike

## Abstract

Sexual reproduction in yeast typically occurs under nutrient-limited conditions, producing stress-resistant spores that promote survival. However, wild *Schizosaccharomyces japonicus* strains isolated from fruit flies exhibit constitutive sporulation even under nutrient-rich conditions. Here we demonstrate that this nutrient-independent sporulation enhances survival by enabling spores to resist digestion by *Drosophila melanogaster*, and that although such strains are disadvantageous under nutrient-rich conditions, they are positively selected during insect predation. Genetic analyses reveal that multiple mutations in six meiotic regulatory genes, acting via epistasis, underlie this phenotype. This trait is widely distributed across natural populations throughout Japan. These findings suggest that constitutive sexual reproduction functions as an adaptive strategy facilitating insect-mediated dispersal, reflecting an ecological context where sexual reproduction is maintained despite its costs. This work provides new insights into yeast–insect symbiosis, evolutionary dynamics of sexual adaptation, and potential applications in yeast breeding.

## Introduction

Sexual reproduction is generally considered less advantageous than asexual reproduction due to its higher costs in time and energy, even in unicellular organisms such as yeast^1,2^. While yeast produces isogamous gametes—gametes of equal size—the classical ‘twofold cost of sex’ (in which sexual populations grow at half the rate of asexual ones because they produce males) does not strictly apply^3^, sexual reproduction nevertheless imposes substantial costs^1^. Consequently, yeasts have evolved tightly regulated mechanisms that restrict entry into the sexual cycle to specific environmental cues—yet the question remains as to why sexual reproduction is maintained in these organisms^4–7^.

Under nutrient-rich conditions, haploid fission yeast proliferates asexually, whereas deteriorating nutrient availability induces sexual differentiation and, ultimately, spore formation^4^. Nitrogen starvation is a key trigger for meiosis between h^+^ and h^−^mating-type cells^8,9^. First, repeated adhesion events occur between the two cell types, forming a multicellular sexual aggregate. This is followed by conjugation between the two cell types, progression through meiosis, and eventual spore formation to resist external stress. Upon sensing nitrogen limitation, the master transcription factor Ste11 activates a transcriptional cascade that promotes meiosis and sporulation^10^. Because this developmental transition is irreversible, activation of *ste11* is tightly regulated^4–7^. Sporulation, which involves meiosis, may represent an adaptive strategy for escaping nutrient-poor environments and colonizing new habitats, as meiosis can generate genetic diversity through increased mutation rates^3,11–13^.

Yeasts form mutualistic associations with insects that facilitate yeast dispersal by transporting spores to new environments^14^. While it is well established that non-motile fungi rely on sexually produced, stress-resistant propagules for survival and dispersal, the extent to which long-standing yeast–insect interactions have driven adaptive evolution, rather than simply reflecting exaptation (the use of a trait for a function it did not originally evolve for), remains unclear^15^. Many insect species consume yeasts and frequently harbour them in their gut or on their body surfaces^16–19^, while insects gain nutritional benefits from specific yeast and bacterial species, indicating reciprocal associations^20–23^. Additionally, insect passaging experiments can select for increased spore production^24^. However, whether such ecological relationships have driven stable evolutionary adaptations in yeast remains unknown^15,25^.

The fission yeast species now known as *Schizosaccharomyces japonicus* was first isolated from a strawberry in Fukuoka in 1928^26–28^. This isolate became the type strain and is widely distributed across Japan. Under nutrient-rich conditions, the type strain rarely forms asci; however, nitrogen starvation rapidly induces sexual reproduction, resulting in asci containing eight spores—a phenotype known as h^90^, which denotes strains capable of homothallic (self-fertile) mating and sporulation^29,30^.

Recently, wild *S. japonicus* strains isolated from fruit flies in Japan were found to undergo frequent sporulation via meiosis even under nutrient-rich conditions—a trait referred to as constitutive sporulation^31^. Previously, such nutrient-independent sporulation was observed only in laboratory-generated mutants^32–37^. The discovery of this trait in natural populations challenges the established view that sexual reproduction in yeast is strictly regulated by nutrient availability and suggests that constitutive sporulation may provide an adaptive advantage in certain ecological contexts. Given the association of these strains with fruit flies, we hypothesised that constitutive sporulation represents an adaptive response to insect-mediated ecological pressures and sought to determine its evolutionary significance in natural yeast populations. Our experimental results confirm that strains possessing the trait of constitutive sporulation under nutrient-rich conditions constitute an alternative adaptive strategy in natural populations.

## Results

### Sporulation in wild *S. japonicus* strains during the log phase

In addition to the eight wild *S. japonicus* strains previously reported^31^, eight newly isolated *S. japonicus* strains from wild fruit flies were examined for their ability to form spores under varying nutrient conditions. Of these, 11 wild strains exhibited high-frequency sporulation on nutrient-rich YE medium in liquid cultures and on agarose plates (**Figure S1**). This phenotype—constitutive sporulation under nutrient-rich conditions—is hereafter referred to as hc^90^.

Upon inoculation of the hc^90^ strains—comprising a mixture of yeast cells and spores—into fresh YE liquid medium, spores germinated and yeast-form cells predominated during the early log phase. As the cultures progressed through this phase, cells aggregated and formed zygotes. Spores within the zygotes matured and were then released following ascus breakdown (**Figure 1a-d**). In contrast, the type strain NIG2021 exhibited little to no sexual aggregation and rarely produced spores under the same conditions (**Figure 1e, f**). In the wild hc^90^ strains (TN33, TN83, and TN84), sexual aggregations were detectable in the early log phase, as indicated by fluctuations in optical density measurements (**Figure 1g**); by the mid-log phase, nearly all cells were aggregated (**Figure 1h**).

**Figure 1.**
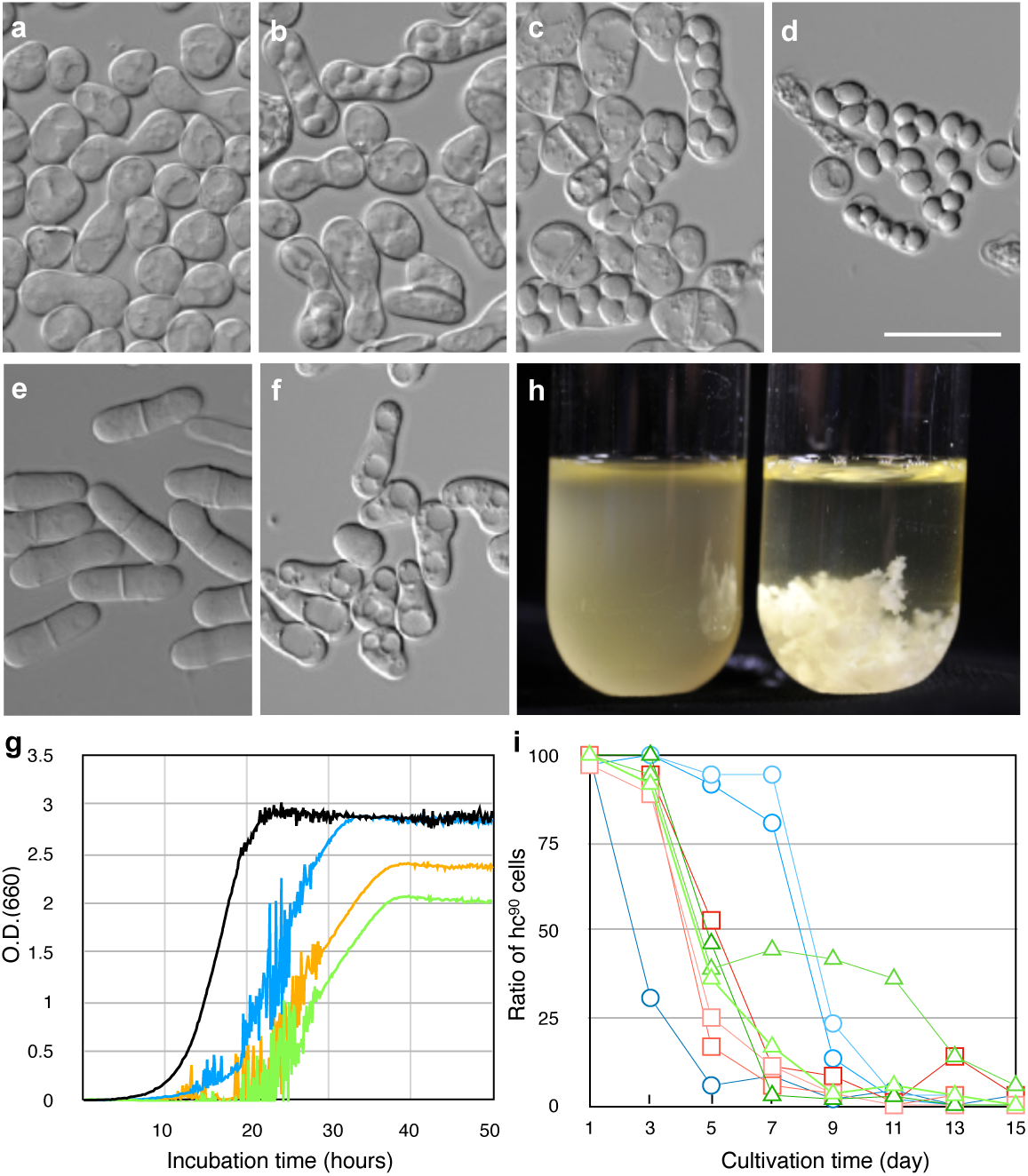
Sporulation and transition of the population under nutrient-rich conditions. **a**-**d**, Representative microscopic images of TN33: **a-d**, (**a**) yeast and zygotes in early log phase, (**b**) immature asci in mid log phase, (**c**) mature asci in late log phase, (**d**) spores released from an ascus during stationary phase. **e-f**, Microscopic images of NIG2021: (**e**) log phase, (**f**) stationary phase. Scale bar, 20 μm. **g**, Cell proliferation monitored by optical density (OD): NIG2021 (black), TN33 (green), TN83 (blue), TN84 (yellow). **h**, Sexual aggregation during log-phase culture in test tubes: NIG2021 (left), TN33 (right). **i**, Serial passaging of TN33 (red), TN83 (green), and TN84 (blue); three independent colonies per strain.

TN40 was previously reported as h^90 31^ (**Figure S1**). Upon careful re-examination of this strain, we determined that it actually exhibits the hc^90^ phenotype; the initial classification as h^90^ was due to the emergence and predominance of sporulation-defective suppressor mutants. Despite the nitrogen-rich conditions of the medium, these wild strains consistently entered the sexual cycle and sporulated extensively. In contrast, four wild strains that failed to sporulate on YE medium were examined in EMM medium lacking ammonium chloride, a standard condition for inducing sporulation (**Figure S1**). These four strains did not produce spores under either condition. Cultures contained only vegetative yeast cells and no zygotes, indicating that the strains were likely sterile.

### Loss of sexual reproduction during serial passaging

Since asexual reproduction is generally more efficient for rapid population expansion under nutrient-rich conditions, sexually reproducing populations may undergo evolutionary shifts during prolonged cultivation. To monitor temporal dynamics within these populations, single clones of the hc^90^ wild strains (TN33, TN83, TN84) were inoculated into YE liquid medium and serially passaged 15 times (**Figure 1i**). The ratio of sexually reproducing cells to vegetative cells in nutrient-rich medium gradually decreased during continuous cultivation, ultimately leading to the disappearance of sexually reproducing cells. In the mid-passage cultures, sexual aggregations were no longer observed, and by the final passage, nearly all cells were vegetative. Although the timing of this decline varied among the strains, the proportion of hc^90^ clones within the yeast population decreased drastically across all cultures, falling below 5% by the final passage. These observations suggest that the original hc^90^ clones were replaced by suppressor mutants that had lost the ability to form spores under nutrient-rich conditions, and that these suppressor mutants emerged and/or were selected during serial passaging.

### Resistance to digestion by a fruit fly

Spores of *Saccharomyces cerevisiae* and *Schizosaccharomyces pombe* are known to resist digestion by fruit flies (*Drosophila melanogaster*)^19^. To assess whether this is also true for *S. japonicus*, hc^90^ wild strains were fed to *D. melanogaster* as a dietary supplement, and the viable yeast cells in the resulting faeces were quantified (**Figure 2a**). No live yeast cells were recovered from the heterothallic strain (h□), which is unable to produce spores unless it mates with a compatible partner (h□), whereas the homothallic strain (capable of self-mating and sporulation) persisted through digestion by *D. melanogaster*. The hc^90^ strains exhibited even greater survival than the homothallic strain. A significant number of spores were observed in the faecal suspensions from the hc^90^ strains, while those from the other strains showed only digestive debris (**Figure 2b**). Furthermore, yeast colonies derived from the faeces of the homothallic strain were able to grow even after treatment with 30% ethanol, which primarily kills vegetative yeast cells (**Figure S2**). These results indicate that *S. japonicus*, like budding yeasts, possesses resistance to digestion by the fruit fly.

**Figure 2.**
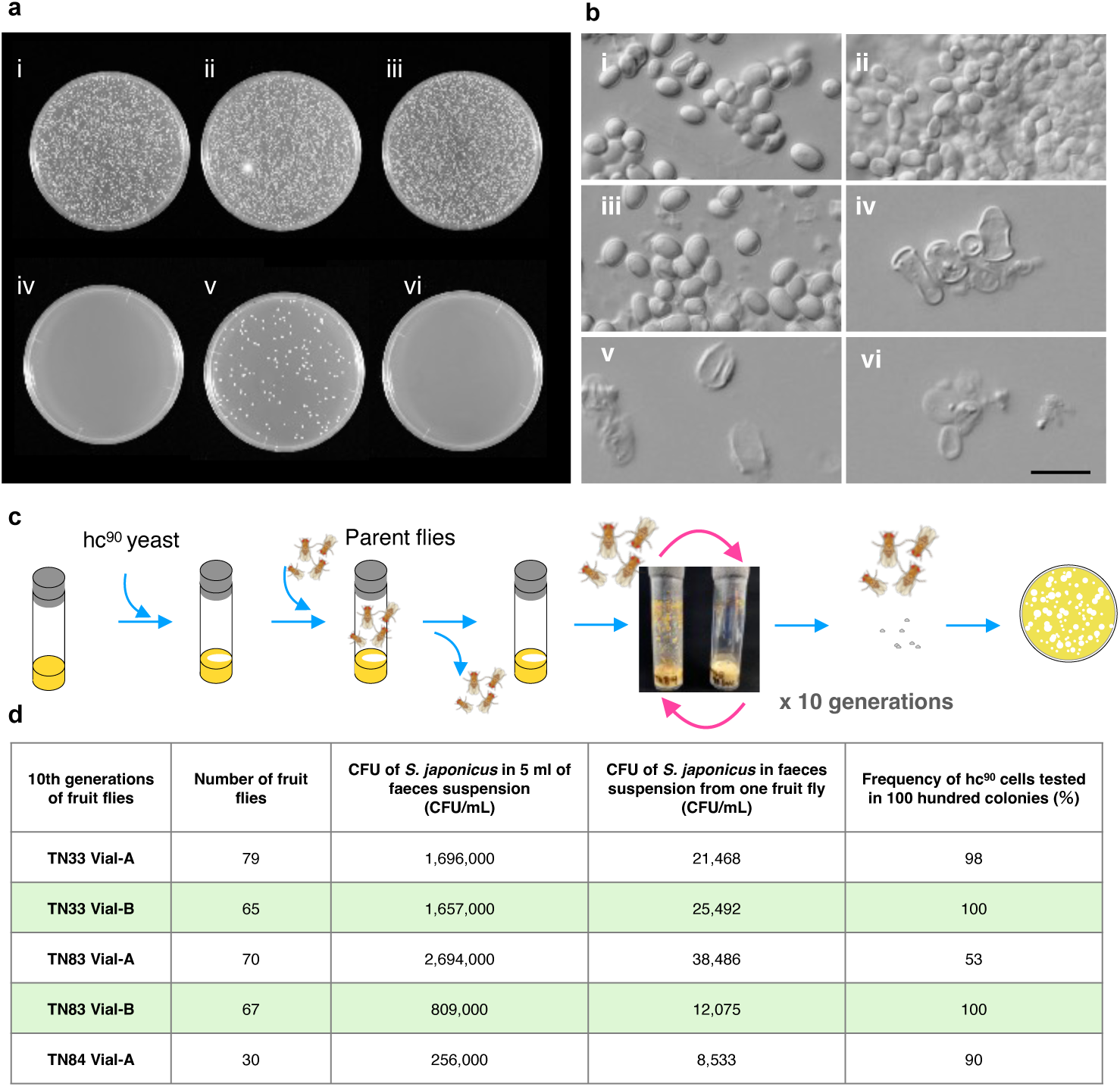
Yeast in faeces and serial passaging via fruit flies. **i**: TN33 (hc^90^); **ii**: TN83 (hc^90^); **iii**: TN84 (hc^90^); **iv**: NIG2017 (h^−^); **v**: NIG2021 (h^90^); **vi**: no inoculation (control). **a,** Agar plates with fly faecal suspensions. Panels i - **v**: 10^−1^ dilution; **vi**: undiluted suspension. **b**, Microscopic images of faecal suspension. Scale bar, 20 μm. **c**, Serial passage schematic for hc^90^ strains via flies. hc^90^ strains were inoculated into fly breeding vials. Parental flies were introduced into these vials, where yeast sporulated. Each fly generation takes approximately 10 d at 25°C. Yeast was transmitted to the new vials by the flies. Serial passage was repeated for ten generations. **d**. Frequency of hc^90^ cells in faeces of tenth-generation flies. Yeast colonies were isolated, and the frequency of hc^90^ colonies in the population was assessed. Two vials per strain were incubated during the tenth generation; however, one TN84 vial was lost due to contamination.

### Serial passage culture of yeast by a fruit fly

To investigate whether insect-mediated ecological pressures drive the persistence and transmission of hc^90^ wild strains, co-culture experiments used yeast and the fruit fly *D. melanogaster*. hc^90^ strains were inoculated into fly breeding vials (**Figure 2c**). The parental flies were kept in the vials for 2 days before removal. After larvae hatched and reached adulthood (first generation), they were transferred to fresh vials, allowing yeast transfer exclusively via flies. While some yeast may have been passively transmitted on fly body surfaces, the primary mode of transfer was likely spore-containing faeces. Yeast colonies were subsequently isolated from tenth-generation fly faeces, and hc^90^ frequency was assessed by scoring sporulation phenotypes (**Figure 2c**). Remarkably, hc^90^ clones remained consistently high even after 10 serial passages. In contrast to laboratory serial passaging, where hc^90^ clones were rapidly eliminated (**Figure 1e**), fly-mediated serial passage maintained a high frequency of hc^90^ clones. These results suggest that fruit flies can exert positive selective pressures on high-sporulation strains even when such strains are lost during laboratory cultivation.

### Selection of the hc^90^ yeast by a fruit fly

To confirm the positive selection of the hc^90^ yeast by a fruit fly, we tested whether feeding a mixed yeast population with a reduced proportion of hc^90^ cells would increase their relative abundance. Full-growth cultures from the serial passage experiment (**Figure 1e**) were maintained at −80°C as glycerol stocks. Glycerol stocks prepared from the eighth-passage cultures were inoculated into fresh YE medium. An aliquot of recultivated cells was plated on YE agar to determine the proportion of hc^90^ yeast in the population, while another aliquot was inoculated into vials for fly breeding. Fruit flies were introduced into the vials and kept for 2 days, after which they were transferred to empty vials to collect faeces. The faeces suspension was plated on YE agar, and the frequency of hc^90^ colonies was determined among the grown yeast (**Figure 3a**).

**Figure 3.**
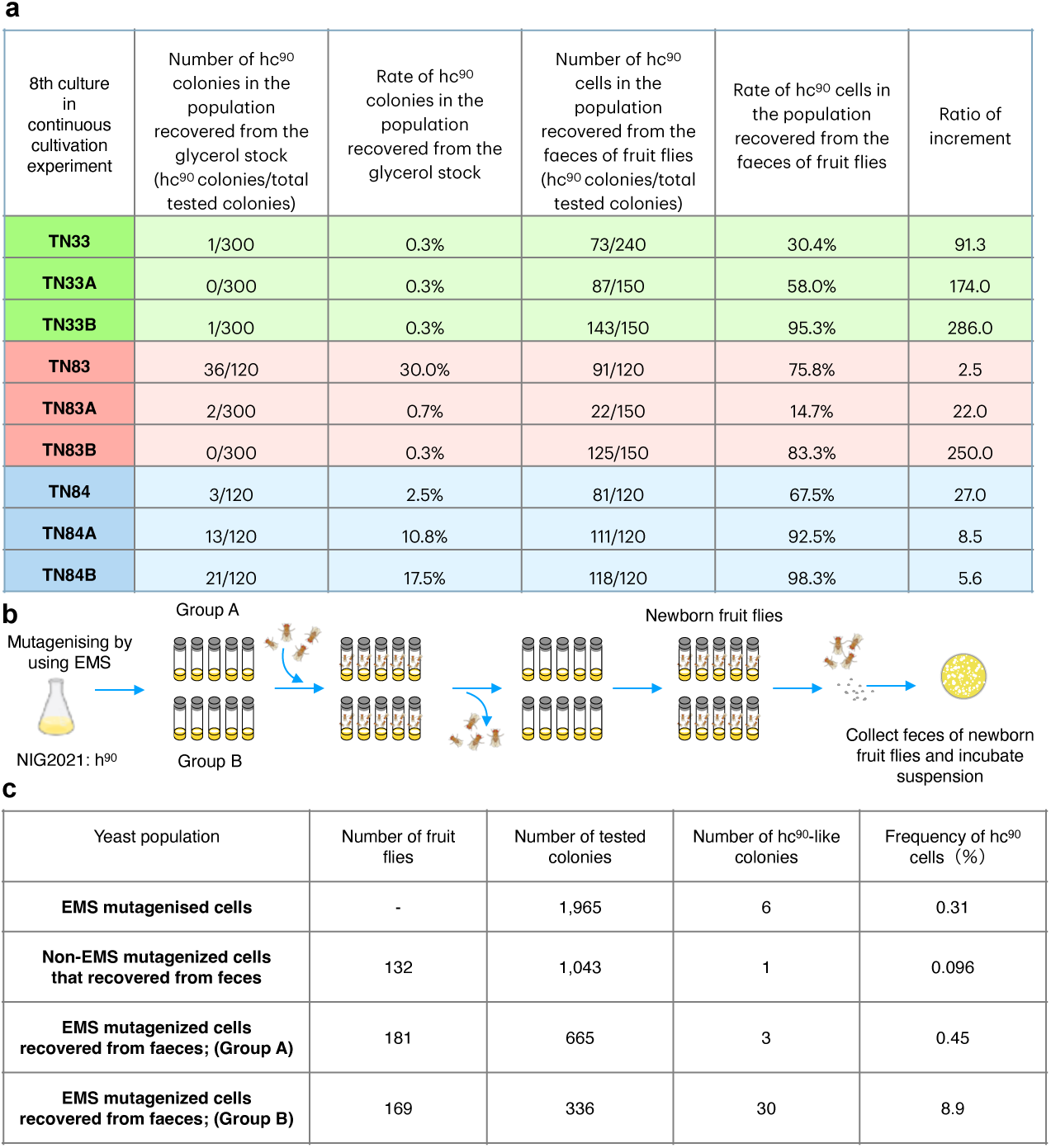
Selection of hc^90^-like yeast by flies from serial passage populations and a chemically mutagenized yeast library. **a**, Eighth-passage cultures under nutrient-rich conditions were preserved as frozen stocks. Each stock culture was inoculated into fly breeding vials, and flies were added. After 2 days of incubation, faeces were collected from the flies, and the frequency of hc^90^ colonies in the faeces was assessed. **b**, Selection schematic for hc^90^-like strains from a chemically mutagenized yeast library. Either mutagenized yeast or non-mutagenized yeast was inoculated into fly breeding vials. Five vials constituted one unit: one experimental unit received non-mutagenized yeast (control), while two units received mutagenized yeast (Groups A and B). **c**, Frequency of hc^90^ colonies in faeces after mutagenized or non-mutagenized yeast feeding. Yeast colonies were isolated from faeces, and the frequency of hc^90^ colonies in each population was assessed.

In two of the serial passage cultures, no hc^90^ colonies were detected in 300 examined colonies from the glycerol stock. The highest frequency observed before fly breeding was 30%. However, in all cases, the frequency of hc^90^ yeast increased after exposure to flies. A particularly dramatic increase was observed in the TN33B culture, where the frequency surged from approximately 0.3% to 95.3%. The results demonstrate that *S. japonicus* strains exhibiting the hc^90^ phenotype can increase in frequency after passage through fruit flies, which act as ecological filters via digestion and subsequent faecal deposition and recovery. Thus, fly-mediated predation may serve as a strong selective pressure favoring the hc^90^ phenotype under laboratory conditions.

### Biological selection of hc^90^-like yeast in a mutant library

If our hypothesis is correct, the hc90 trait may have arisen naturally as a result of positive selective pressure exerted by fruit fly predation, initially selecting for spontaneous mutants displaying nutrient-independent sporulation. To test this hypothesis, we sought to isolate hc^90^-like mutants capable of sporulating under nutrient-rich conditions from a yeast mutant library (**Figure 3b**).

The type strain NIG2021 was mutagenized with ethyl methanesulfonate (EMS) to generate a mutant library. Screening among 1,965 colonies identified 6 clones capable of sporulating in YE medium, representing 0.31% of the total population (**Figure 3c**). Yeast from this library was then fed to adult flies; after laying eggs, the parents were removed and the newly emerged adults (the next generation) were collected. Faeces from these flies were then used to recover yeast cells.

Two replicate experimental groups (Groups A and B) were established (**Figure 3a-b**). In Group A, three YE-sporulating clones were obtained from 665 colonies (0.45%), a frequency not significantly different from that in the original mutant library. In contrast, Group B yielded 30 YE-sporulating clones from 336 colonies, representing a notable increase in frequency to 8.9%. All recovered clones produced spores in YE medium, although the fraction of sporulating cells in the population was lower than that observed in hc^90^ wild strains. The results suggest that the fruit fly’s ability to digest yeast cells while preserving spores within its faeces constitutes a natural selection mechanism that can enrich for yeast strains with enhanced sporulation traits.

### Genetic variations in sporulation-related genes of hc^90^ strains

To identify genes associated with the hc^90^ trait, we analysed single-nucleotide polymorphisms (SNPs) in three hc^90^ strains. Whole-genome sequencing revealed numerous SNPs in the three fruit fly–derived strains, TN33, TN83, and TN84, compared with the type strain^38,39^ :47,725 in TN33, 58,970 in TN83, and 47,439 in TN84. These strains shared 17,013 SNPs, which made it difficult to identify those linked specifically to the hc^90^ phenotype. Accordingly, we focused on SNPs in genes related to meiosis and sporulation.

In the fission yeast *S. pombe*, Ste11 functions as the master transcription factor that drives the transition from asexual to sexual reproduction (**Figure 4a**). It regulates the expression of approximately 80 genes involved in meiosis and sporulation^6^. The expression of *ste11* is tightly controlled by the TORC1 pathway, the stress-responsive MAPK pathway, and cAMP signalling pathway, while the mating pheromone–responsive MAPK pathway activates its transcription. Orthologs of these regulatory components are conserved in *S. japonicus*^40^, and orthologs of *ste6, byr2, win1*, and *ste11* are known to be essential for sporulation^41^. We therefore searched for amino acid substitutions in the meiosis- and sporulation-related factors represented in Figure 4a. Six genes—*byr2*, *mcs4*, *ste6*, *ste11*, *tor2*, and *win1*—were found to carry amino acid substitutions in all three strains (TN33, TN83, and TN84) **(Figure S3**). Some of these mutations were shared among the three strains, while others were strain specific.

**Figure 4.**
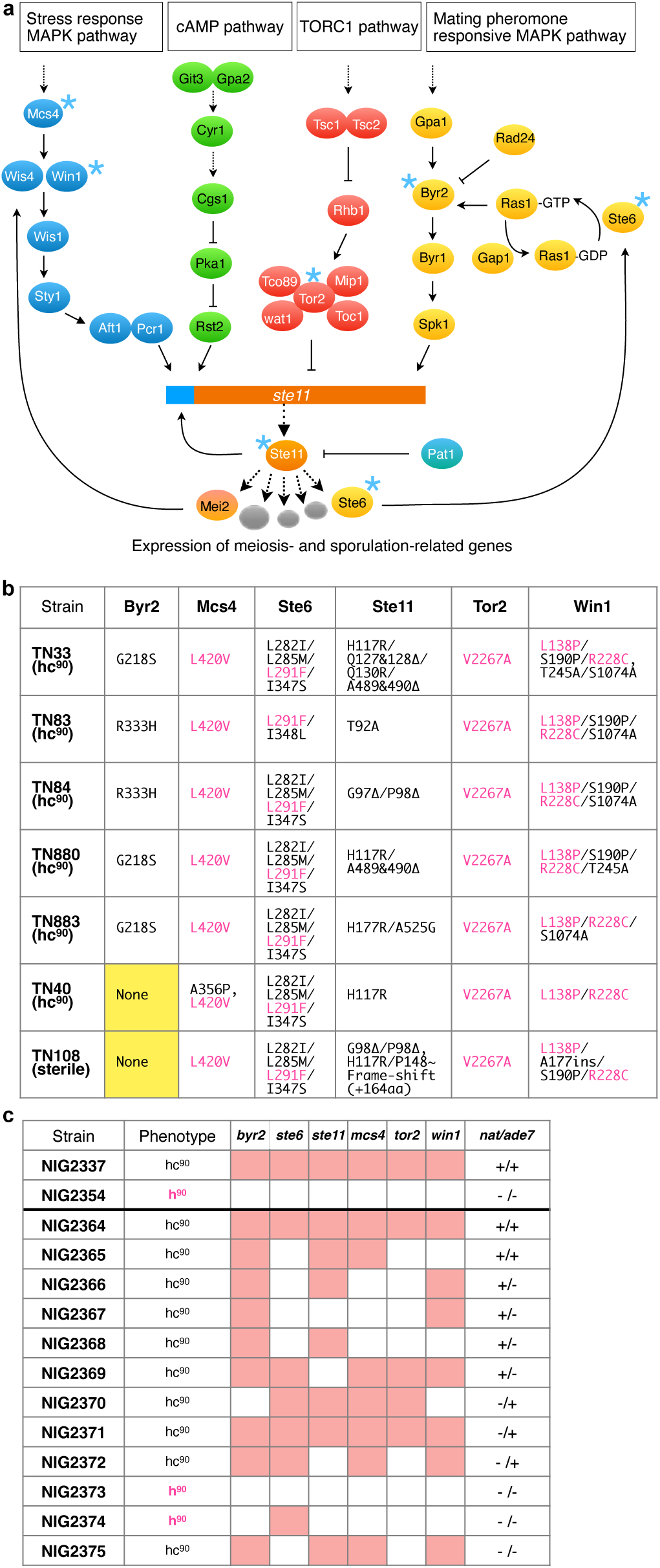
Genetic variations in sporulation-related genes of hc^90^ strains. **a**, Signalling network regulating meiosis and sporulation^8–10,55^. Asterisks indicate proteins with amino acid variations commonly found in fly-isolated hc^90^ strains. **b**, Amino acid sequence variations in gametogenesis regulatory elements (Byr2, Mcs4, Ste6, Ste11, Tor2, and Win1) associated with the hc^90^ phenotype. **c**, Genetic analysis of progeny from a cross between NIG2337 (a TN33 derivative, hc^90^, *nat*^+^, *ade*7^+^) and NIG2354 (h^90^, *nat^−^*, *ade*7^−^). The hc^90^ phenotype is shown in magenta, and mutations of *byr2*, *mcs4*, *ste6*, *ste11*, *tor2*, and *win1* are shown in pale red boxes.

### Genetic cross of hc^90^ strains

To investigate the genetic basis of the hc^90^ trait, we performed a genetic cross using TN33 derivative NIG2337 (*natMX*, nourseothricin-resistant). This strain was crossed with an h^90^ strain carrying *ade7* (NIG2354). Progeny were categorised based on nourseothricin resistance and/or the *ade7* mutation, enabling efficient identification of hybrid spores. We then analysed the hc^90^ phenotype and the genotypes of the six candidate regulatory genes (**Figure 4b**). The results indicated that none of the six genes alone was sufficient to confer the hc^90^ trait, although not all combinations of mutated alleles were recovered in progeny.

### Suppressor mutants arising during serial passage

We analysed suppressor mutants that emerged during serial passaging lost the ability to form spores on YE medium. A total of 52 suppressor mutants from the eight-passage cultures were tested for their ability to sporulate under nutrient-poor conditions. This analysis revealed a clear distinction between fertile (19 suppressors) and sterile (33 suppressors) phenotypes. The 19 fertile suppressors were classified as revertants, resembling the type strain, which undergoes sexual reproduction in a nutrient-dependent manner.

We focused on the *ste11* gene because it was previously reported to suppress uncontrolled meiosis and sporulation^10^, and we identified novel mutations in 16 of these suppressor mutants (**Figure 5**). One revertant carried a frameshift mutation downstream in *ste11*, resulting in a truncated protein that nonetheless restored Ste11’s function to a level comparable to that of the type strain despite retaining the original mutations (P98&G97Δ, H117R) (**Figure 5b**). In contrast, one sterile suppressor harboured a truncated gene product due to a nonsense mutation, while 12 carried frameshift mutations producing altered proteins (**Figure 5c**). Two additional sterile suppressors exhibited amino acid substitutions within the HMG (**h**igh-**m**obility **g**roup) box, an essential DNA-binding domain of transcription factors^42^. The gene products of these sterile suppressors lacked either the functional HMG box or a major portion of the Ste11 protein, resulting in the sterile phenotype.

**Figure 5.**
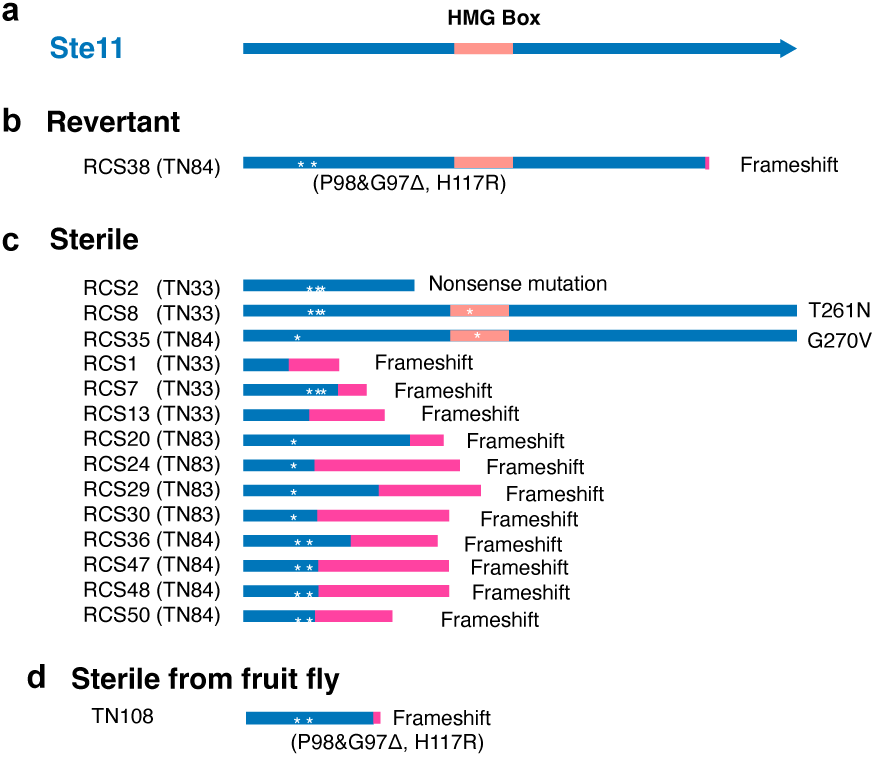
Suppressor mutations in *ste11*. **a**, Ste11 full-length sequence (624 amino acids), including the HMG box. **b**, Revertant mutant derived from TN84 that exhibits nutrient-dependent sporulation, like the type strain. **c**, Sterile suppressors derived from TN33, TN83, and TN84. **d**, Frameshift mutation in fly-isolated strain TN108. Asterisks indicate amino acid substitutions, and additional amino acids from the frameshift are shown in magenta.

These suppressor mutations, which arose spontaneously during serial passage under non-mutagenized conditions, suggest that constitutive sporulation on YE medium can be suppressed by naturally occurring mutations, even in the absence of intentionally elevated mutation rates. Notably, a similar mutation was found in a sterile wild strain (TN108) isolated from a fruit fly (**Figure 5d**).

### Constitutive sporulation in wild *S. japonicus* isolated from diverse sources

To determine whether the hc^90^ phenotype of *S. japonicus* is exclusive to strains isolated from fruit flies, wild strains were collected from various plant and soil sources (dandelion flowers, periwinkle, morning glory, moss, and soil) across northern, central, and southern Japan (**Figure 6a-d**). Dandelion flowers were frequently visited by insects (**Figure 6e & 6f**), and one strain was isolated from honey bees on goldenrod (*Solidago virgaurea* var*. asiatica*) (**Figure 6g & 6h**). In total, 22 strains were collected. Except for one strain isolated from morning glory (NIG2229), all displayed the hc^90^ phenotype, showing frequent sporulation even on YE medium.

**Figure 6.**
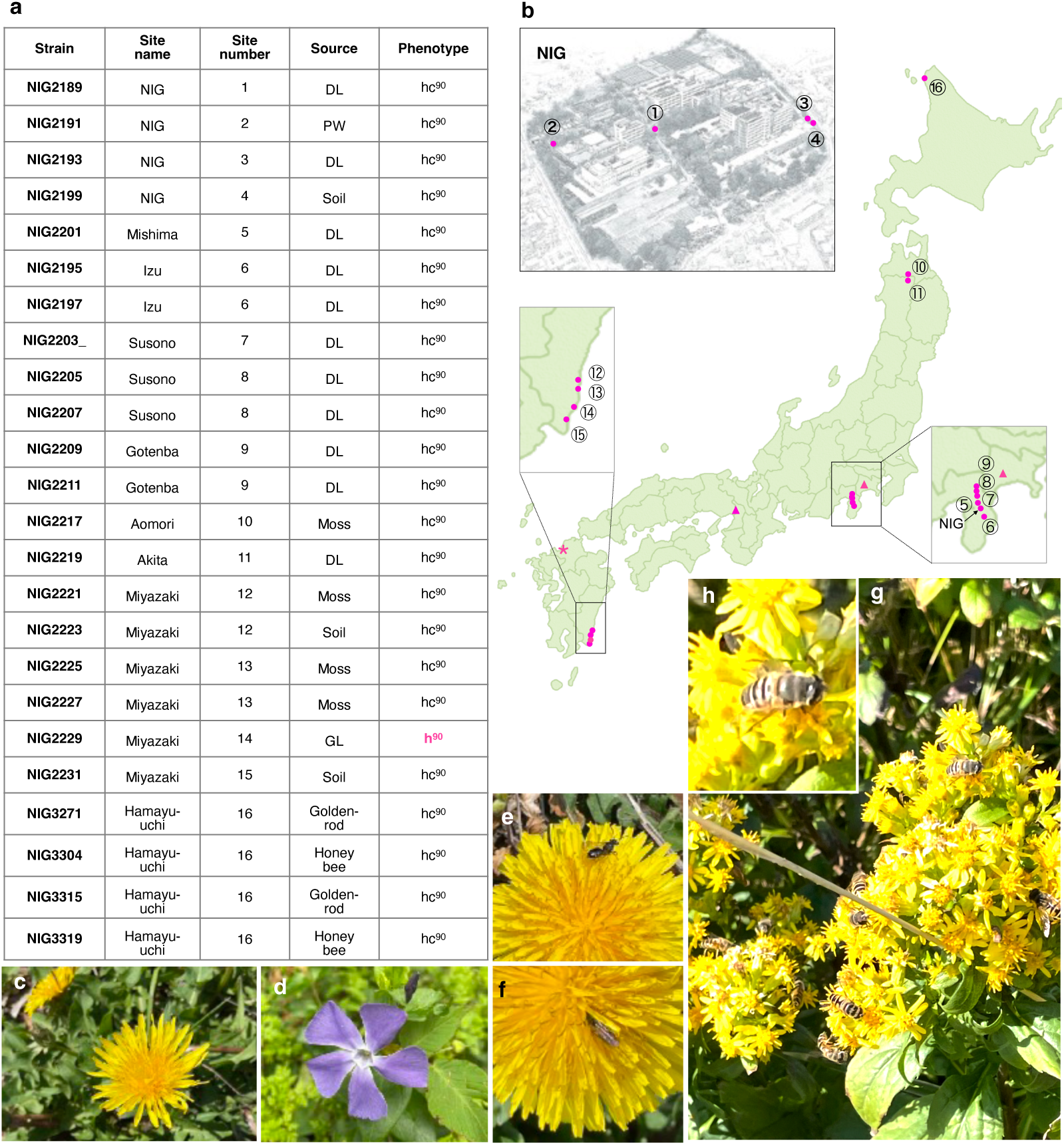
Isolation of wild *S. japonicus* yeast. **a**, Wild *S. japonicus* strains from Japan. DL: dandelion; PW: periwinkle; GL: morning glory. **b**, Collection sites in Japan. Triangles indicate sites where yeast was isolated using banana traps for flies, and circles indicate direct isolation from natural sources. An asterisk marks the site where the type strain was originally isolated in 1928. Numbers correspond to panel a. Aerial view of the National Institute of Genetics (NIG) is also shown. **c**, Dandelion. **d**, Periwinkle. **e–f**, Insects on dandelions. **g–h**, Honeybees on goldenrod with magnified view.

We analyzed mutations in six genes: *byr2*, *mcs4*, *ste6*, *ste11*, *tor2*, and *win1* (**Figure S4**). No amino acid substitutions were found in NIG2229. In contrast, the other strains carried amino acid substitutions in these genes, although one hc^90^ strain (NIG2231) lacked mutations in *byr2*. Notably, genetically distinct hc^90^ variants occurred within a narrow geographical area: the four hc^90^ wild strains isolated from different sources within our institute exhibited unique combinations of these variants (**Figure 6b**). These findings indicate that *S. japonicus* strains with the hc^90^ phenotype, and with diverse genetic backgrounds, are widely distributed across Japan’s natural environments, inhabiting not only fruit flies but also flowers, mosses, and soil. The isolation of an hc^90^ wild strain from honey bees further supports the role of insects as vectors for transferring *S. japonicus* between flowers. Insects were also frequently observed visiting dandelion flowers.

### Other wild yeasts with high-frequency spore production

Some wine yeast strains have been reported to sporulate even on nutrient-rich media^43,44^. To examine whether this phenomenon extends to other budding yeasts, we examined the sporulation capacity of wild *Saccharomyces cerevisiae* strains in our yeast collection. In total, 11 strains of *S. cerevisiae* were isolated from diverse environments, including fruit flies, grapes, moss, and soil. Laboratory strains of *S. cerevisiae* are generally unable to sporulate on nutrient-rich media such as YPD, standard budding-yeast medium^45^, yet 10 of the wild isolates sporulated on YPD medium (**Figure S5a-d**). These wild *S. cerevisiae* strains proliferated as vegetative cells during both log and stationary phases, with sporulation increasing sharply after 3 days in stationary-phase culture. This sporulation pattern was observed in both liquid and solid (agar) media, although its timing differed from the pattern seen in *S. japonicus*.

Budding yeast *S. cerevisiae* possesses regulatory genes for gametogenesis that are functionally analogous to those in fission yeast^46^, and nitrogen starvation combined with the absence of a fermentable carbon source induces the expression of the master regulator of gametogenesis, *IME1*^4^. After *IME1* is expressed, it induces the secondary transcription factor *NDT80*, irreversibly committing the cells to meiosis. Mutations in *IME1* and/or *NDT80* were identified in a subset of these isolates, though not in all (**Figure S5 e**). Our findings raise the possibility that fruit flies may positively select yeast species with nutrient-independent sporulation, such as *S. japonicus*.

## Discussion

Our findings suggest that meiosis and sporulation under nutrient-rich conditions—a hallmark of the unstable hc^90^ phenotype—may represent an adaptive strategy for survival in insect-associated environments. We hypothesise that, in such environments, insect hosts selectively digest vegetative cells, whereas spores remain intact and thereby enable persistence and transmission. This interpretation is supported by model co-culture experiments under controlled conditions, in which yeast strains with the hc^90^ phenotype coexisted with fruit flies while maintaining sporulation activity. Moreover, hc^90^ strains are widely distributed across Japan and exhibit substantial genetic diversity in *ste11* and its upstream regulatory elements, indicating that this phenotype persists across populations. Collectively, our results provide evidence that positive selection by insects can enrich for yeast strains with constitutive sporulation in our experimental system. Our co-culture system using yeast and fruit flies may model the selection and development of industrially relevant yeast strains.

Although six genes were identified as candidates associated with the hc^90^ phenotype, none alone was sufficient to confer this trait (**Figure 4c**). Combinations of at least two mutations among these genes appear necessary for the hc^90^ phenotype. The strong association of these six genes with the hc^90^ phenotype is supported by 23 wild hc^90^ strains isolated from independent sources carrying genetic variations in the same six genes (**Figure S4**). However, not all possible combinations of the six mutated genes were recovered in progeny (**Figure 4c**), suggesting additional genes may be involved. Furthermore, a revertant suppressor (TN108) with *ste11* mutations strongly suggests that the mutations leading to the hc^90^ phenotype occur in genes that regulate Ste11 function (**Figure 5c**). Thus, these six genes, including *ste11* and its upstream regulatory genes, are likely key genetic determinants of the hc^90^ phenotype.

Experimentally, mutations that enable nutrient-rich sporulation have been isolated in *cyr1*, *pat1*(*ran1-3*), *pka1*, and *rad24* mutants^32–34,36,37^. If positive selection by insects acted solely on spore survival, evolution of meiosis-independent sporulation would also be advantageous. Indeed, a *pat1* mutant showing meiosis-independent sporulation under nutrient-rich conditions has been isolated^33,34^. However, no genetic diversity has been observed in *rad24*, *pka1*, or *pat1* among the wild isolates in this study. These mutants exhibit impaired vegetative growth and reduced spore germination rates, likely making them unfit for survival in natural environments^33,34,36,37^. The cAMP signalling pathway is involved in a wide range of cellular regulatory processes, making it unlikely to produce the hc^90^ phenotype via this pathway alone^47^. Similarly, the TORC1 and stress-responsive MAPK pathways are unlikely to act alone. Nevertheless, if these pathways contribute to the hc^90^ phenotype, each mutation is likely to exert only subtle functional effects on the phenotype, and the phenotype may result from epistatic interactions among multiple mutations. This interpretation is consistent with genetic crosses and wild hc^90^ strains possessing mutations in nearly all candidate genes for this phenotype.

Nearly all wild hc^90^ strains retain mutations in all six genes. Loss of the diverse combinations of these variations—including partial and complete loss—could arise through sexual reproduction because the six genes are dispersed across three chromosomes (**Figure**lJ**S3b**). However, inter-strain outbreeding among hc^90^ strains may act as a buffer against the erosion of these genetic combinations. Moreover, as fruit flies provide an environment that promotes outbreeding in yeast^48^, such outbreeding may greatly expand combinatorial variant diversity, producing epistatic effects that contribute to the hc^90^ phenotype^49^. This suggests that fruit flies, beyond selecting digestion-resistant spores, may play a role in maintaining genetic combinations for the hc^90^ phenotype.

The replacement of hc^90^ yeast by suppressors emerging during serial passaging is likely because, under favourable nutrient conditions, asexual reproduction is far more advantageous for increasing cell numbers than sexual reproduction. When two cells mate and undergo meiosis, they form a set of eight spores, whereas a vegetative cell divides six or seven times in the same period, producing approximately 32 or 64 progeny cells. Furthermore, spores require time to revert to vegetative cells before resuming cell division. Under strong positive selective pressure against sexual reproduction, populations are expected to shift rapidly from sexual to asexual reproduction. Nevertheless, the frequency of suppressor emergence appears to be relatively high. Because *S. japonicus* does not respond to iodine vapour staining—a standard method for assessing sporulation efficiency^26^—direct quantification of suppressor frequency is not possible. Instead, in *S. pombe*, where sporulation-defective mutants such as *ran1-3* (*pat1*) can form spores under nutrient-rich conditions without undergoing meiosis, the emergence frequency of such mutants has been estimated at roughly one in every 50,000 cells^33^. Thus, suppressor mutations that abolish constitutive sporulation appear to arise at a relatively high frequency in model yeasts such as *S. pombe*^50^. Although the underlying mechanisms remain unclear, one plausible explanation is that the mutation rate increases during meiosis^13^, contributing to the observed frequency of suppressor emergence.

To our knowledge, this is the first successful isolation of *S. japonicus* from the northernmost part of Japan. Yeast strains exhibiting the hc^90^ phenotype were found across Japan, suggesting that, while human activities may have contributed to their dispersal, insects could also play a significant role in their geographic distribution. The detection of similar combinations of genetic variations among isolates from diverse sources implies that the yeast may have circulated between insects and plants, expanding its range through insect-mediated transmission.

Attempts to isolate this yeast from other insects, such as ants collected from flowers, have thus far been unsuccessful. To date, successful isolations have been achieved only from fruit flies and honey bees. Winged insects were frequently observed visiting sampled flowers, suggesting a potential ecological link between flowers and the insects carrying the yeast (**Figure 6e-h**). The yeast was also frequently isolated from moss, a major source of isolates; however, there is currently no direct evidence that it actually grows within moss tissues. It remains to be determined whether the moss environment merely provides conditions favourable for the long-term survival of spores.

Only one *S. japonicus* strain exhibiting the h^90^ phenotype—sporulation in response to nutrient conditions—has been isolated from natural environments, namely from a morning glory flower (NIG2229). The wild strain shared the same six genotypes as the type strain, with no amino acid substitutions detected. These findings suggest that there may be natural environments in which the h^90^ phenotype, characterised by nutrient-dependent sporulation, is strongly favoured. However, most of the isolates obtained from field surveys were hc^90^-type yeasts, and the primary habitats of h^90^ yeasts remain unknown. Notably, one sterile strain isolated from a fruit fly carried a frameshift mutation in the *ste11* gene, similar to the suppressor mutants experimentally obtained in the laboratory (**Figure 5d**). This observation implies that, in nature, hc^90^ yeasts may also have encountered conditions in which asexual reproduction was more advantageous, as observed in the laboratory serial passage experiment. Alternatively, it is possible that hc^90^ strains may have a competitive advantage over natural h^90^ yeast populations in the regions surveyed so far.

Our results raise fundamental questions about adaptive evolution. Interactions with insects appear to shape the survival, evolution, and adaptation of yeast strains with the hc^90^ trait in natural environments. Consequently, genetic populations of hc^90^ may be distinct from h^90^, and indeed, many hc^90^ strains have been isolated from diverse regions across Japan. In contrast, h^90^ phenotypes are rarely found, raising the question of what kinds of environments might favor this trait and where such environments could exist.

Yeast has evolved a sophisticated system for sensing nutrient availability—particularly nitrogen sources—to regulate entry into the reproductive cycle. Nevertheless, evidence from this work suggests that some yeast populations may override this regulatory system in favour of an alternative adaptive strategy. Why would such a strategy evolve? If this phenomenon is indeed driven by interactions with insects, it is notable that yeasts and insects have maintained such associations for over 100 million years^51–53^. Given the duration and ecological intimacy of these associations, it seems likely that adaptive evolution, such as the hc^90^ trait, has occurred during this extended period. Further symbiotic experiments between yeast and insects will be essential for understanding the evolutionary and ecological dynamics underlying their natural interactions.

## Methods

### Yeast cultivation

Yeast strains were cultured in YE medium (Bacto^TM^ yeast extract 5 g/L, glucose 30 g/L) and Edinburgh minimal medium 2 (EMM2)^54^. Agar was added to prepare plate medium (Bacto^TM^ agar 20 g/L). For sporulation tests, nitrogen-free EMM2 medium (lacking ammonium chloride) and Bacto^TM^ malt extract (Becton, Dickinson and Company, MD, USA) were used. All cultures were incubated at 30°C. Antibiotics were used at the following concentrations: G418 (40 μg/mL), nourseothricin (Nat) (100 μg/mL), and aureobasidin A (0.5 μg/mL).

### Yeast strains

The type strain of *Schizosaccharomyces japonicus* used in this study was NIG2021^30^, derived from IFO1609, the original strain isolated by Yukawa and Maki (1931). The strain NIG2354 (h□□, *ade7*; nonsense mutation at W126) was derived from NIG2021. NIG2337 (hc□□, *natMX*), derived from TN33, was constructed by introducing a DNA fragment containing *ste11* and the *natMX* marker gene into TN33. The *S. japonicus* strains used in this study are available from the National Institute of Genetics Bioresource Center via JapoNet (https://shigen.nig.ac.jp/yeast/japonet/).

### Monitoring of optical density

Cell growth and sexual aggregation were monitored using a rocking incubator equipped with an optical density sensor (TVS062CA, ADVANTEC, Tokyo, Japan). An L-shaped culture vial containing 5 mL of cell suspension was incubated with gentle shaking (30 rpm) at 30 °C. Optical density at 660 nm (OD□□□) was measured at 5-min intervals, and vials were allowed to stand for 30 s before each measurement. The total monitoring period was 66 h.

### Isolation of wild yeast

Plant and soil samples collected from natural environments were placed in 50 mL conical tubes. An appropriate volume of YE medium containing antibiotics was added. To inhibit bacterial growth, antibiotics included ampicillin (100 μg/mL), kanamycin (20 μg/mL), chloramphenicol (30 μg/mL), and tetracycline (30 μg/mL). To inhibit fungi, aureobasidin A (0.5 μg/mL) was added to the medium, as *S. japonicus* is resistant to this agent. To isolate *S. japonicus*, samples were incubated at 37°C for 2 days, as *S. japonicus* is resistant to high temperatures. After yeast growth, cultures were spread on YE agar plates to obtain single colonies, which were then re-streaked onto fresh YE plates. Yeast cell morphology was observed under a microscope to confirm the presence of *S. japonicus*. The *S. japonicus* strains used in this study are available from the National Institute of Genetics Bioresource Center via JapoNet (https://shigen.nig.ac.jp/yeast/japonet/).

### Fruit fly cultivation

The wild-type *Drosophila melanogaster* Oregon-R strain was obtained from NBRP Drosophila (https://nbrp.jp/en/resource-search-en/). Fly maintenance and egg laying were conducted at 25°C. The fly food medium was prepared by suspending dried yeast powder Y2A (Asahi Group Foods, Ltd., Japan) at 32 g/200 mL and, separately, suspending agar (0.8 g) and glucose (40 g) in 200 mL. Both mixtures were autoclaved at 120°C for 20 min and combined on a hot stirrer at 150°C. Cornmeal powder (16 g) (Oriental Yeast Co., Japan) was added, followed by stirring for an additional 20 min. After reducing the temperature to 100°C, antibiotics (ampicillin 100 μg/mL, kanamycin 20 μg/mL, chloramphenicol 30 μg/mL, tetracycline 30 μg/mL) and aureobasidin A (0.5 μg/mL) were added. Approximately 8 mL of the prepared medium was dispensed into sterile transparent plastic vials and sealed with sponge plugs. All medium preparation and dispensing were performed inside a clean cabinet. Completed fly food vials were stored at 4°C.

### Sporulation test

To test yeast sporulation, 0.1–0.2 mL of overnight yeast culture was inoculated into fly food vials and incubated at 30°C for 2 d. Approximately 30 adult flies were then added and the vials were incubated at 25°C for an additional 2 d. For faeces collection, flies were transferred into 50 mL conical tubes and maintained at 25°C for 1–2 d. Flies were removed, and 5 mL of sterile water was added and vortexed to generate a faecal suspension. This suspension was diluted and spread on YE agar plates, then incubated at 30°C to detect yeast.

The cells were washed with sterile water and resuspended in 1 mL of 30% ethanol, then incubated at 37 °C for 30 min to kill any remaining vegetative cells. The surviving spores were washed with sterile water and resuspended in 1 mL of sterile water. This spore suspension was spread on YE agar plates and incubated at 30 °C for 2 d to allow colony formation.

### Coexistence experiment

For yeast–fly coexistence studies, 0.1–0.2 mL of overnight yeast culture was inoculated into fly food vials and incubated at 30°C for 2 d. Approximately 30 adult flies were then introduced and reared at 25°C for 2–3 d to allow egg laying, after which all adult flies were removed. Emerged adult flies were transferred to fresh food vials. To prevent the transfer of the original yeast-containing medium, the mouths of the old and new vials were joined with the new vial positioned above, and the paired vials were placed upright on a bench. After 5 min, most flies naturally moved upward. The vials were then sealed with sponge plugs. This process was repeated to maintain successive generations.

### Yeast mutagenesis

A single colony of NIG2021 was inoculated into YE medium and cultured overnight at 30 °C. An aliquot (0.3 mL) of the overnight culture was then inoculated into 60 mL of YE medium and incubated for an additional overnight period at 30 °C. The overnight culture was centrifuged (1,000 rpm, 5 min) to harvest the cells; the supernatant was discarded, and the pellet was resuspended in 30 mL of EMM medium. The suspension was aliquoted (9 mL each) into separate tubes, and 99% ethyl methanesulfonate (EMS; Sigma-Aldrich, USA) was added at 45, 90, or 140 µL. Cells resuspended in the remaining EMM medium without EMS were used as a control. The cultures were incubated at 30 °C for 3 h. Subsequently, 5 mL of 5% sodium thiosulfate was added to each culture, followed by an additional 40 min of incubation. Cells were then washed twice with EMM medium and finally resuspended in 1 mL of EMM. Aliquots of 100 µL were dispensed and stored at −80 °C. After freezing, viable cell numbers were quantified as colony-forming units for both the EMS-treated and control samples. Samples showing approximately a 10% survival rate after EMS treatment were used as the EMS-treated library.

### Genetic cross

NIG2354 (h□□, *ade7*; nonsense mutation at W126) was pre-cultured in 3 mL YE medium and incubated overnight at 30 °C. NIG2337 (hc□□, *natMX*) was pre-cultured in 100 mL YE medium and incubated overnight at 30 °C. For mating, the NIG2337 culture was used at an early growth stage when all spores had germinated and only yeast-form cells were present. The NIG2354 culture was centrifuged and resuspended in 1 mL nitrogen-free EMM2 medium. An aliquot of 0.3 mL of this suspension was added to 5 mL of nitrogen-free EMM2 medium. The NIG2337 culture was likewise centrifuged and resuspended in 1 mL nitrogen-free EMM2 medium. After measuring optical density, NIG2337 cells (less than one-tenth the NIG2354 concentration) were added to the medium containing NIG2354. The mixed culture was incubated statically at 30 °C overnight. After confirming completion of sporulation under a microscope the following day, cells were harvested and resuspended in 1 mg/mL Zymolyase (Funakoshi, Tokyo, Japan) solution and incubated at 37 °C for 1 h. Cells were washed with sterile water and resuspended in 1 mL 30% ethanol, then incubated at 37 °C for 30 min to kill remaining vegetative cells. Spores were washed with sterile water and resuspended in 1 mL sterile water. This spore suspension was spread on YE agar plates and incubated at 30 °C for 2 d to allow colony formation.

### Microscope

Cell images were captured using a Zeiss Axio Imager A2 microscope equipped with a Plan-Apochromat 63×/1.4 oil DIC objective lens (Zeiss). Screening for the hc^90^ phenotype was performed using a Nikon Eclipse Ni-L microscope equipped with a Plan-Fluor 40×/0.75 DIC objective lens (Nikon).

### Gene sequencing

*S. japonicus* strains and 22 *S. cerevisiae* strains were cultured overnight in 3 mL YE medium (*S. japonicus*) or YPD medium (*S. cerevisiae*) at 30 °C with shaking. The following day, cells were harvested, and genomic DNA was extracted using the Wizard® Genomic DNA Purification Kit (Promega, Madison, WI, USA). Prior to DNA extraction, cells were treated with Zymolyase (Funakoshi, Tokyo, Japan) in digestion buffer at 37 °C for 1 h to remove the cell wall. PCR amplification was performed on the extracted genomic DNA using synthetic primers for target genes. *S. japonicus* gene sequences were obtained from JaponicusDB (https://www.japonicusdb.org)^39,40^. Primers used for PCR and sequencing are listed in the Supplementary Excel files.

### Genome sequencing and SNP calling

Genomic DNA from wild *S. japonicus* strains TN33, TN83, and TN84 was sequenced using 150-bp paired-end Illumina technology (NovaSeq platform, Illumina, San Diego, CA, USA) following Illumina DNA PCR-Free Library Prep Kit instructions. All analyses were performed against the reference genome assembly of *S. japonicus* strain SJ5 (GenBank accession GCA_000149845.2).

Raw reads underwent quality control and trimming with fastp (version 0.12.1). Paired-end reads underwent automatic adapter detection, trimming of low-quality bases from the 5′ and 3′ ends, and short-read removal; only reads with a trimmed length > 50 bp were retained. fastp also generated per-sample HTML and JSON quality reports for downstream inspection.

Trimmed reads were aligned to the *S. japonicus* reference genome (GCA_000149845.2_SJ5_genomic.fna) using BWA-MEM (version 0.7.19) with default parameters, including read-group information for each sample. Alignments were sorted and indexed using Samtools (version 1.22.1; https://www.htslib.org). The reference genome was indexed prior to alignment using bwa index and samtools faidx. Single-nucleotide polymorphisms (SNPs) were identified using bcftools (version 1.22). Variants were called assuming a haploid genome, and only high-confidence SNPs were retained, defined as SNP sites with a Phred-scaled quality score ≥30 and a read depth ≥10. Each strain’s final SNP set was normalised against the reference genome and indexed for downstream analyses.

## Acknowledgements

We thank Dr. Jun Kirano (NIG) for critical reading and helpful comments, Dr. Makoto Kawamukai for valuable advice, and Dr. Chikara Hashimoto for insightful discussion. We also thank laboratory technical staff, including Aki Ide, Megumi Kamata, and Masayuki Tachihara, for their assistance. We thank the NBRP *E. coli* and NBRP *Drosophila* for providing strains. We are also grateful to Dr. Kuniaki Saito (NIG) for technical advice on the fly experiments. This research was supported by a “Strategic Research Projects” grant (2023-SRP-09) from ROIS (Research Organization of Information and Systems). This work was supported by JST PRESTO, Japan, Grant Number JPMJPR24N9 to T.S.

## Author contributions

H.N. and T.S. conceived and designed the study. H.N. performed fly–yeast experiments and yeast genetic analyses. H.N., S.N., K.F., T.M., and T.S. conducted gene sequencing experiments. T.S. performed whole-genome sequencing and analysis. H.N. and T.M. constructed yeast strains. S.N., K.F., K.A., T.S., and H.N. conducted field sampling. S.N., T.S., and H.N. performed strain isolation and variant analysis. H.N. analysed the data and prepared all figures. H.N. and T.S. wrote the Methods section. H.N. drafted the manuscript with input from all authors.

## Competing interests

The authors declare no competing interests.

## Declaration of generative AI and AI-assisted technologies in the manuscript preparation process

During the preparation of this work the author(s) used Chat GPT, Grammary, and Nature Research Assistant in order to improve English grammar and clarity. After using this tool/service, the author(s) reviewed and edited the content as needed and take(s) full responsibility for the content of the published article.

---

**Figure S1.**
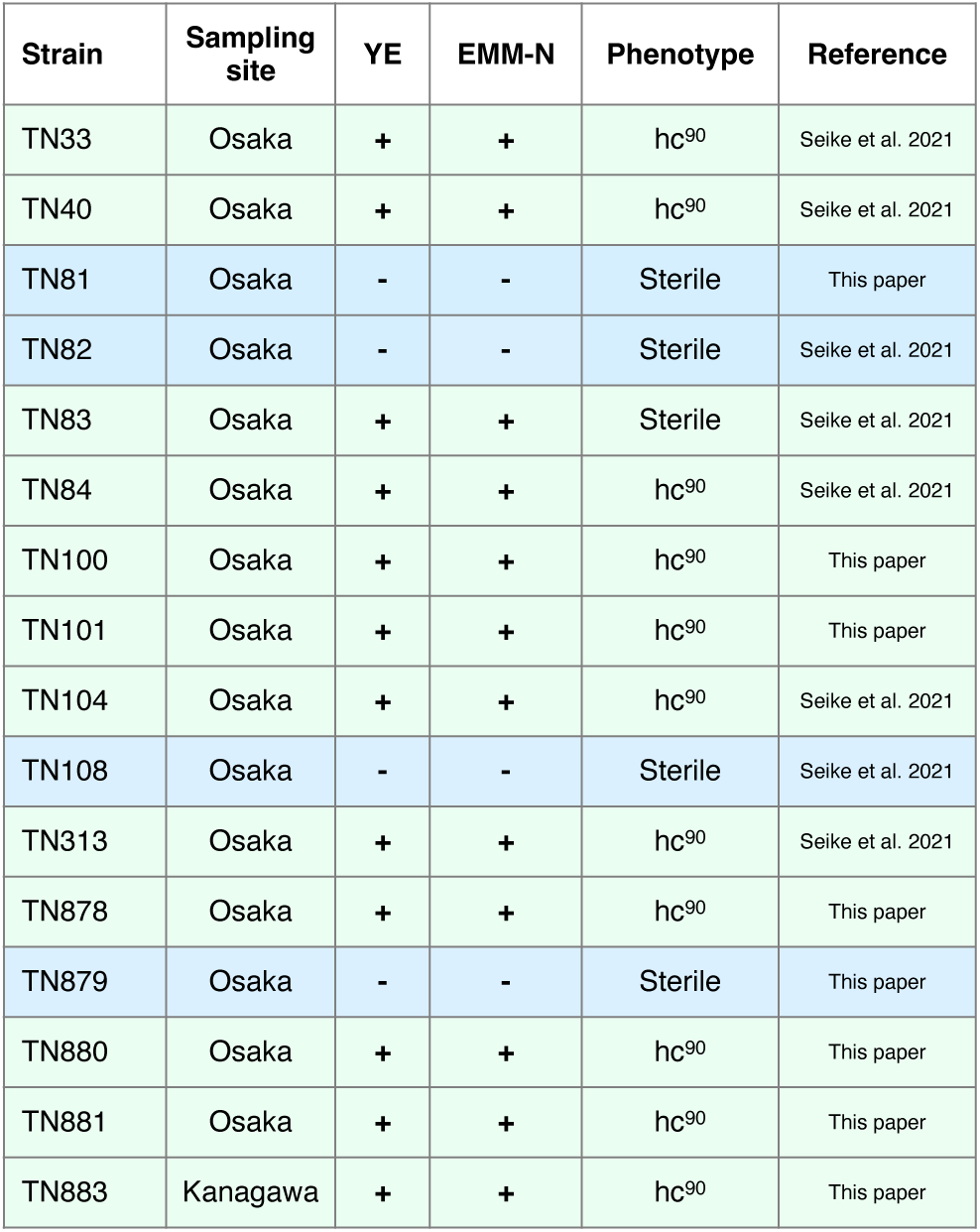
Wild *S. japonicus* strains isolated from fruit flies.

**Figure S2.**
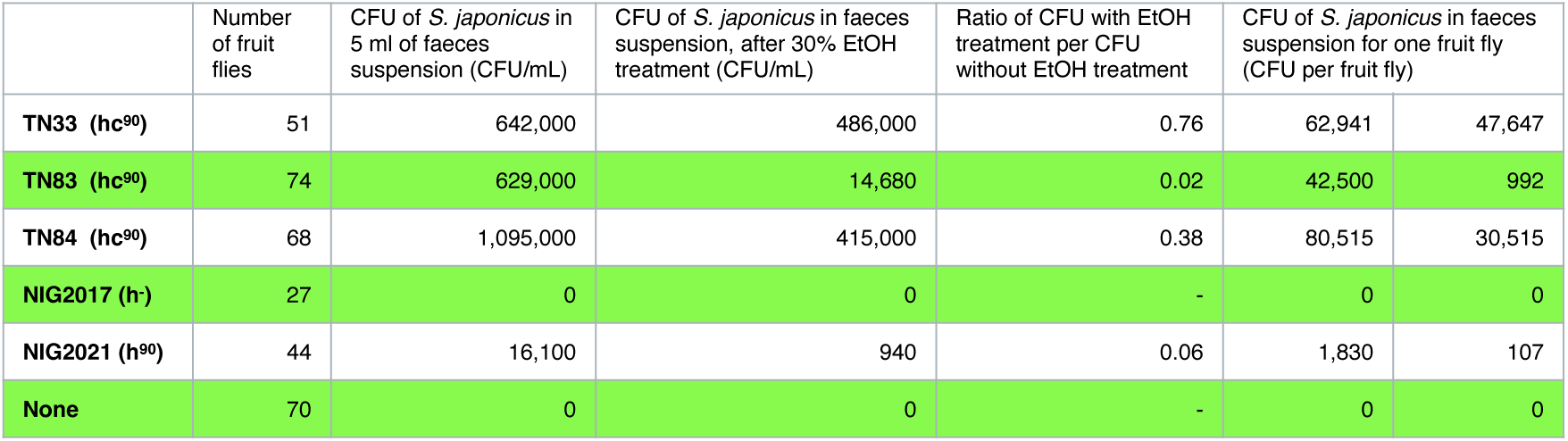
Colony-forming units (CFUs) of *S. japonicus* in fly faeces suspension.

**Figure S3.**
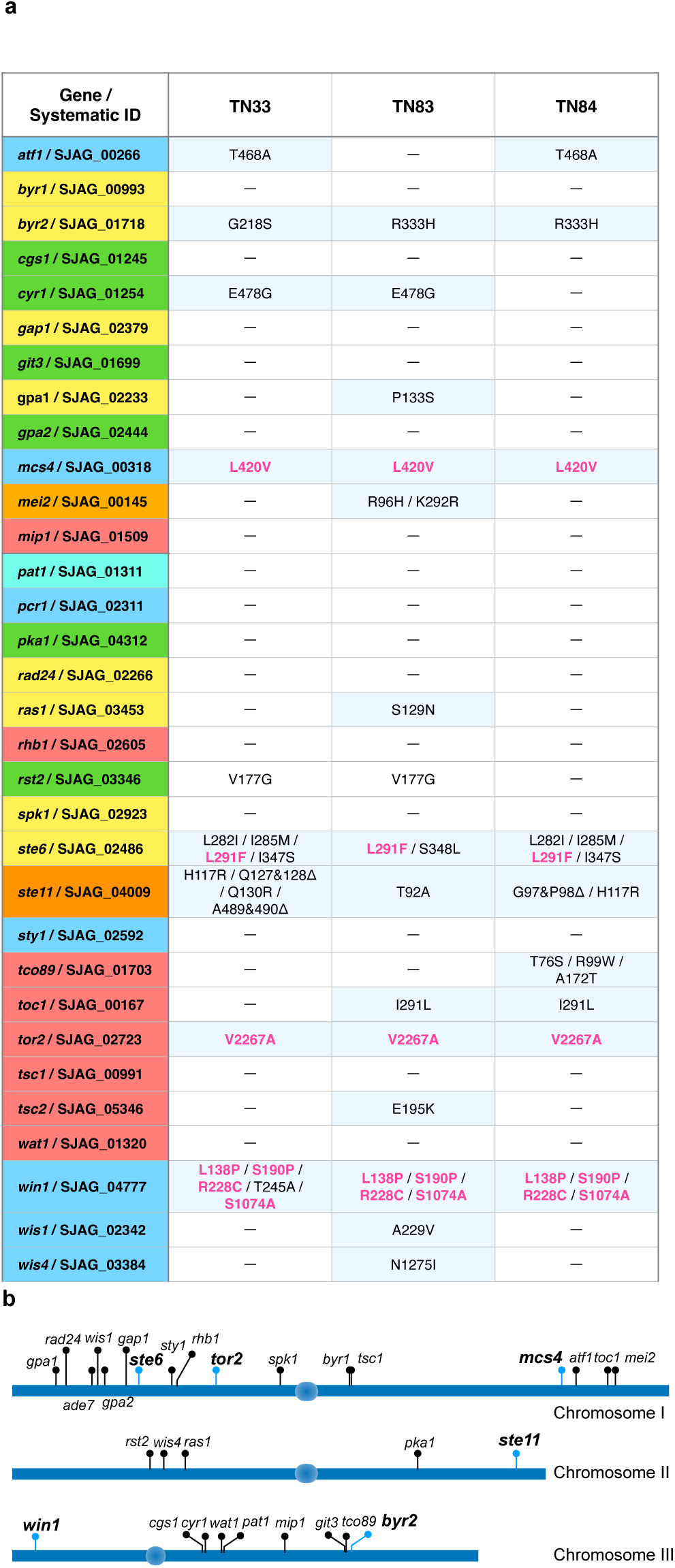
a, Amino acid substitutions in meiotic and sporulation genes. Gene name colors match Figure 4a signaling network elements. Amino acid substitutions shared by TN33, TN83, and TN84 are shown in magenta. **b**, The genetic map of meiotic and sporulation genes.

**Figure S4.**
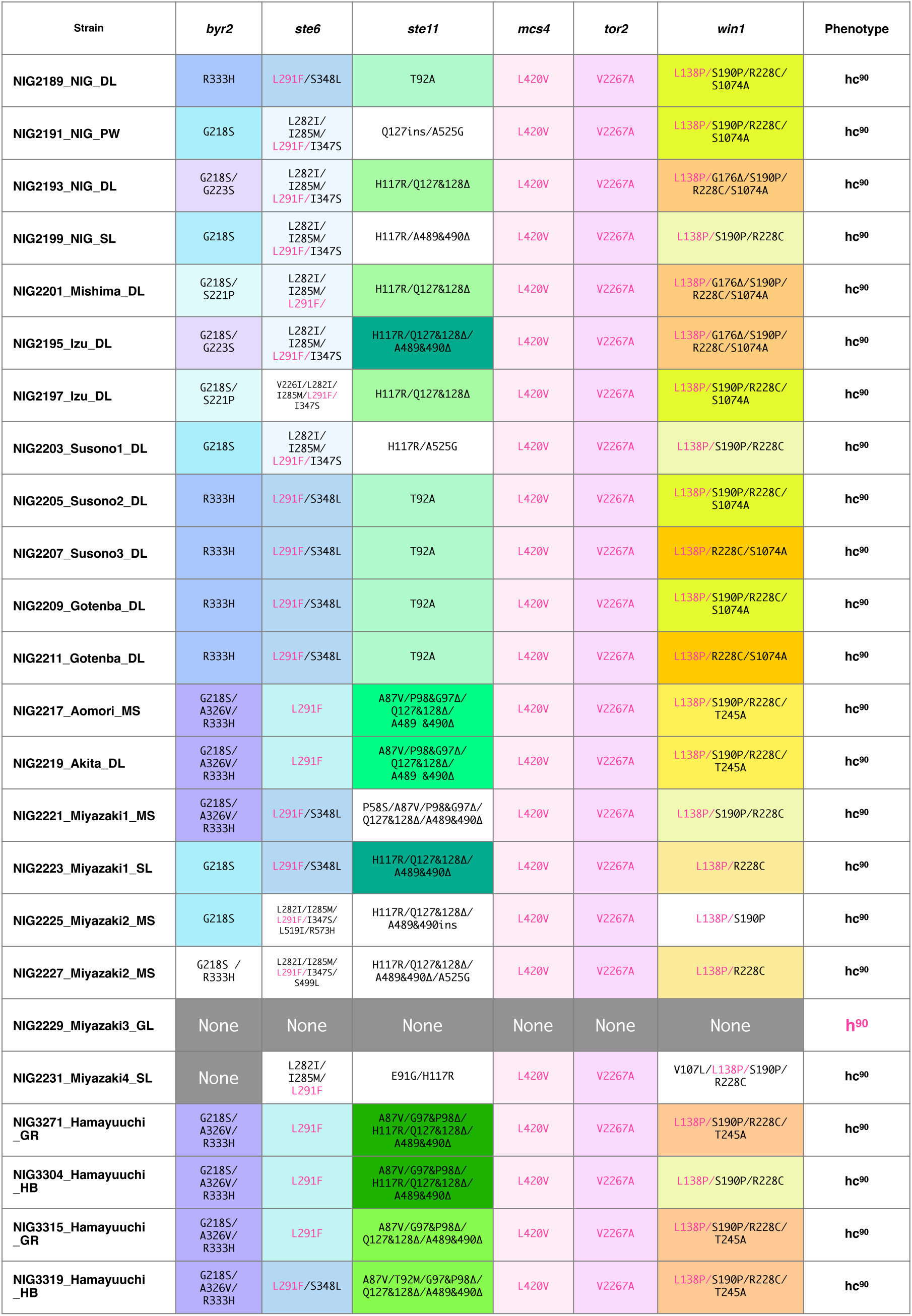
Amino acid substitutions in meiotic and sporulation genes in wild *S. japonicus*.

**Figure S5.**
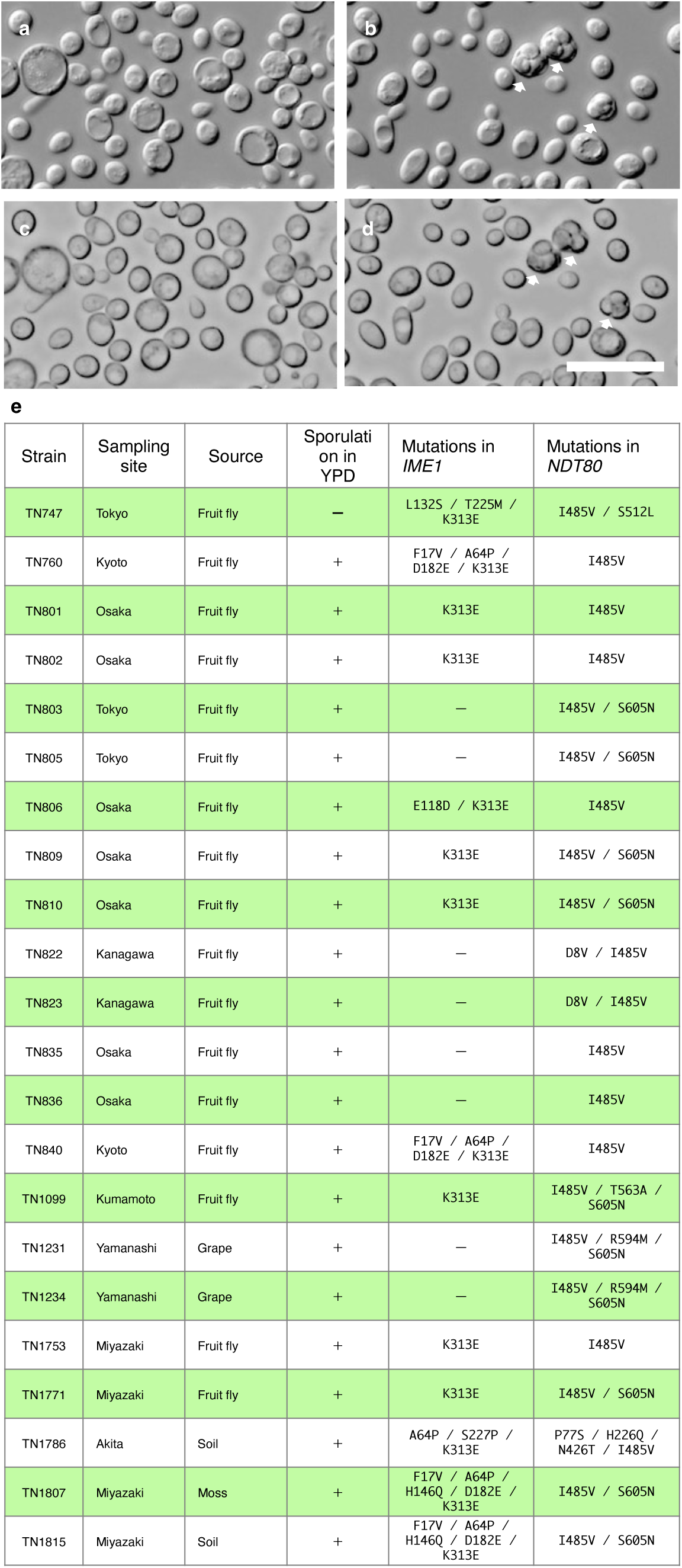
Sporulation of wild *Saccharomyces cerevisiae* under nutrient-rich conditions. **a**, Yeast in YPD medium at day 3. DIC images (upper panels), DIC images with high contrast (lower panels). Arrows indicate asci: W303 (left), TN706 (right). Scale bar, 20 μm. **b**, Wild *Saccharomyces cerevisiae* strains.

## Notes

### Competing Interest Statement

The authors have declared no competing interest.

